# 14-3-3ζ mediates an alternative, non-thermogenic mechanism to reduce heat loss and improve cold tolerance

**DOI:** 10.1101/853184

**Authors:** Kadidia Diallo, Sylvie Dussault, Christophe Noll, Angel F. Lopez, Alain Rivard, André C. Carpentier, Gareth E. Lim

**Affiliations:** Département de médecine, Université de Montréal, Montréal, QC, Canada; Axe cardio-métabolique, Centre de Recherche du Centre Hospitalier de l’Université de Montréal (CRCHUM), Montréal, Québec, Canada; Département de médecine, Université de Sherbrooke, Sherbrooke, QC, Canada; Centre de Recherche du Centre Hospitalier de l’Université de Sherbooke (CHUS), Sherbrooke, QC, Canada; Centre for Cancer Biology, SA Pathology and University of South Australia, Adelaide, South Australia, Australia

**Keywords:** 14-3-3 proteins, 14-3-3ζ, beiging, vasoconstriction, adaptive thermogenesis

## Abstract

Following prolonged cold exposure, adaptive thermogenic pathways are activated to maintain homeothermy, and elevations in body temperature are generally associated with UCP1-dependent and -independent increases in energy expenditure. One of the earliest, identified functions of the molecular scaffold, 14-3-3ζ, was its role in the synthesis of norepinephrine, a key endogenous factor that stimulates thermogenesis. This suggests that 14-3-3ζ may have critical roles in cold-induced thermogenesis. Herein, we report that transgenic over-expression of TAP-14-3-3ζ in mice significantly improved tolerance to prolonged cold. When compared to wildtype controls, TAP mice displayed significantly elevated body temperatures and paradoxical decreases in energy expenditure. No changes in β-adrenergic sensitivity or oxidative metabolism were observed; instead, 14-3-3ζ over-expression significantly decreased thermal conductance via increased peripheral vasoconstriction. These findings suggest 14-3-3ζ mediates alternative, non-thermogenic mechanisms to mitigate heat loss for homeothermy. Our results point to an unexpected role of 14-3-3ζ in the regulation of body temperature.

**Graphical abstract:** 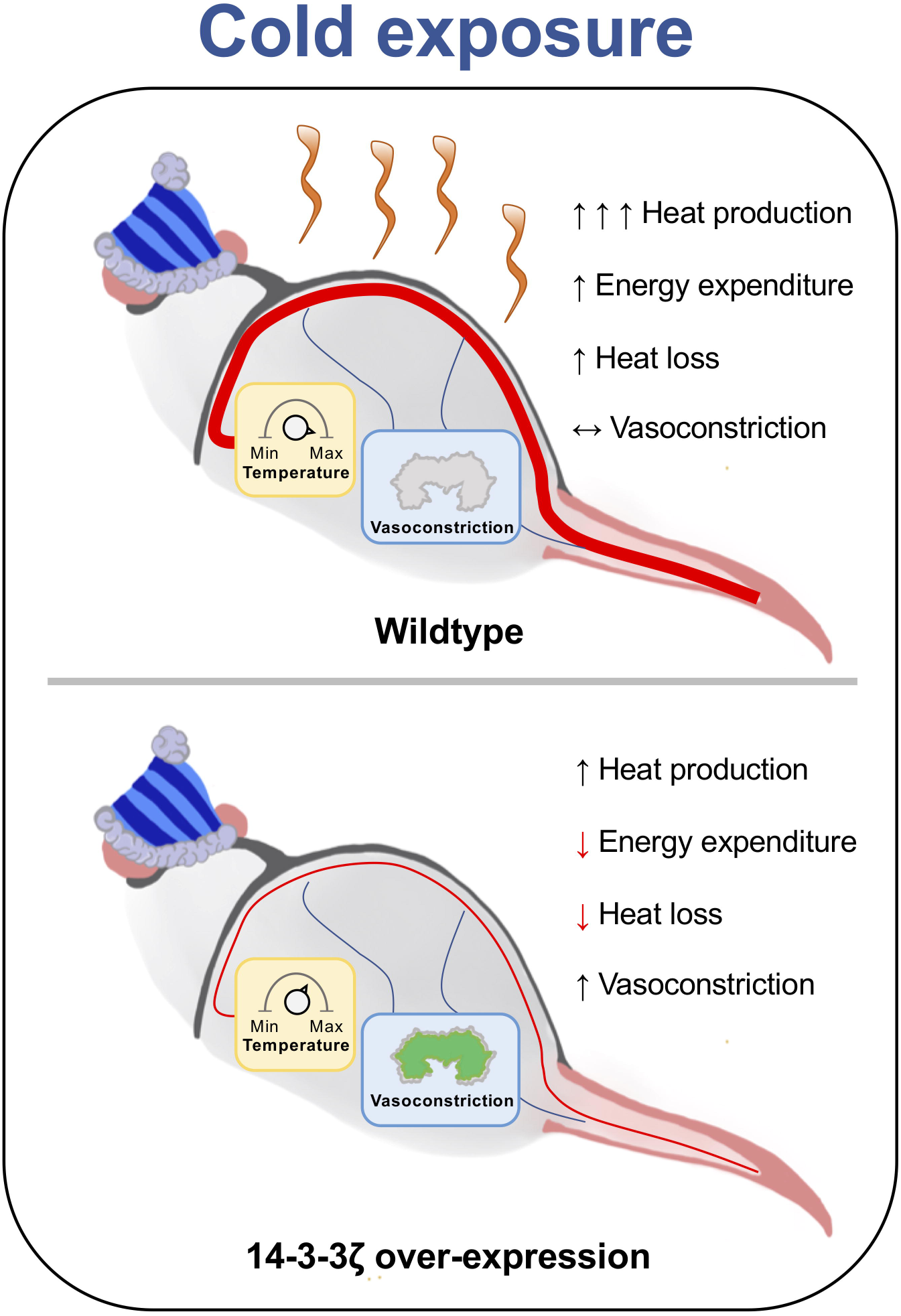

## Introduction

Homeothermy is the maintenance of a stable, internal body temperature following changes in environmental temperature, and it is essential to the survival of endotherms (Morrison and Nakamura, 2019). To defend against hypothermia, mammals have evolved various thermogenic mechanisms, ranging from skeletal muscle shivering to non-shivering, adaptive thermogenesis (Bal et al., 2012; Bal et al., 2017; Morrison and Nakamura, 2019; Palmer and Clegg, 2017; Pant et al., 2016). Shivering, which is short in duration, is the fast contraction of skeletal muscles during which heat is released via the hydrolysis of ATP (Nowack et al., 2017). Another rapid mechanism to regulate body temperature is vasoconstriction, which helps to mitigate heat loss (Wang et al., 2006), but its relative contribution to homeothermy in rodents is often underappreciated.

The sympathetic nervous system (SNS) is the primary regulator of adaptive thermogenesis, as the release of norepinephrine from efferent nerves activates β-adrenergic receptors on the surface of brown adipocytes in rodents to trigger heat production (Cannon and Nedergaard, 2004). Long-term cold exposure can also induce the recruitment of brown-like adipocytes in inguinal white adipose tissue (iWAT) through a process known as beiging (Kajimura et al., 2015; Wang and Seale, 2016). Brown and beige adipocytes are distinct thermogenic fat cells rich in uncoupling protein-1 (UCP1), which uncouples the proton gradient from ATP synthesis to produce heat during fatty acid oxidation (Cannon and Nedergaard, 2004; Kajimura et al., 2015; Wu et al., 2012). Although the contributions of BAT to thermogenesis have been well-established, the relative contributions of beige adipocytes to thermogenesis are unclear (Labbe et al., 2016). However, in the context of an adrenergic stress, such as severe burns, detectable increases in beige adipocyte thermogenic activity have been reported (Salinas Sanchez et al., 1989). Recently, emphasis on investigating the therapeutic potential of treating diabetes and obesity by activating beige and brown adipocytes has been proposed (Barbatelli et al., 2010; Kajimura et al., 2015), but the mechanisms underlying brown and beige adipocytes development and function are still not completely understood.

14-3-3ζ is a member of the 14-3-3 scaffold protein family, which are highly conserved serine and threonine binding proteins present in all eukaryotes (Diallo et al., 2019; Dougherty and Morrison, 2004; Fu et al., 2000; Mugabo et al., 2018). They bind to a diverse number of enzymes, signalling proteins, and transcription factors and have been implicated in the regulation of numerous cellular processes, including proliferation, protein trafficking, and apoptosis (Dougherty and Morrison, 2004; Fu et al., 2000). Recently, we have reported various contributions of 14-3-3ζ to whole-body metabolism, as it was found to be an important regulator of glucose metabolism and adipogenesis (Lim et al., 2015; Lim et al., 2013; Lim et al., 2016). The ability of 14-3-3 proteins to bind to enzymes and exert positive or inhibitory effects on their activities have been documented. For example, interactions between 14-3-3 proteins and RAF-1 or PKA potentiate their kinase activity (Fantl et al., 1994; Kent et al., 2010); in contrast, interactions of DYRK1A with 14-3-3 proteins can attenuate kinase function (Alvarez et al., 2007). Interestingly, one of the first ascribed functions of 14-3-3 proteins is their regulation of tyrosine (TH) and tryptophan (TPH) hydroxylases, both of which are rate-limiting enzymes involved in the synthesis of norepinephrine and serotonin, respectively (Kleppe et al., 2011). Furthermore, norepinephrine and serotonin have been demonstrated to respectively stimulate and inhibit adaptive thermogenesis and beiging (Bartness et al., 2010; Crane et al., 2015; Landsberg et al., 1984; Morrison and Nakamura, 2019). Therefore, given the ability of 14-3-3ζ to regulate the activities of TH and TPH, we hypothesized that 14-3-3ζ may have critical roles in the development and function of beige and brown adipocytes, thereby influencing adaptive thermogenesis.

In the present study, we report on the outcome of reducing or increasing 14-3-3ζ expression in the context of tolerance to acute and prolonged cold stress. Transgenic mice over-expressing 14-3-3ζ were protected from acute and prolonged cold exposure, and in the context of prolonged cold, 14-3-3ζ over-expression was associated with increased body temperature but with a paradoxical decrease in energy expenditure and a lack of differences in oxidative metabolism in BAT and various tissues. Moreover, these effects were not due to changes in the sensitivity of transgenic mice to β-adrenergic stimuli or sympathetic activity. Strikingly, mice over-expressing 14-3-3ζ displayed decreased thermal conductance, or heat loss, due to elevated vasoconstriction. Collectively, our data demonstrate that 14-3-3ζ over-expression improves cold tolerance by activating non-thermogenic mechanisms to mitigate heat loss in the context of cold-tolerance. Furthermore, results from this study highlight the need to consider non-thermogenic mechanisms in the context of cold tolerance or adaptation and caution against invoking solely increased thermogenesis to explain tolerance to prolonged cold exposure.

## Results

### Partial deletion of 14-3-3ζ does not affect acute cold tolerance

To understand whether 14-3-3ζ could influence cold tolerance, we started by examining the effect of reducing or increasing 14-3-3ζ expression in the context of an acute exposure to cold. We previously reported that homozygous 14-3-3ζ knockout mice weighed significantly less than WT mice due to reduced fat mass, which could affect their ability to tolerate cold (Gregory, 1989; Haemmerle et al., 2006; Lim et al., 2015); thus, mice heterozygous (HET) for *Ywhaz*, the gene encoding 14-3-3ζ, were used. WT and HET mice were challenged with acute cold (4°C) for 3 hours, and no differences in body weights were detected between WT and HET mice before and after the acute cold challenge (Figure 1A,B). Furthermore, both groups displayed similar decreases in rectal temperature throughout the entire challenge (Figure 1C). Similar results were observed in female mice (Figures S1A,B). Acute cold exposure significantly increased *Ywhaz* and *Ucp1* mRNA levels in brown adipose tissue (BAT) of WT and HET male mice (Figure 1D, Figure S1C). In inguinal white adipose tissue (iWAT), levels of *Ywhaz* mRNA were significantly increased by cold exposure in WT and TAP mice, but no differences were observed in the expression of *Ucp1* and the beige selective gene *Tmem26* (Figure 1E, Figure S1D), indicating that a 3 hour cold exposure was not sufficient to induce beiging (Kajimura et al., 2015; Wang and Seale, 2016).

**Figure 1.**
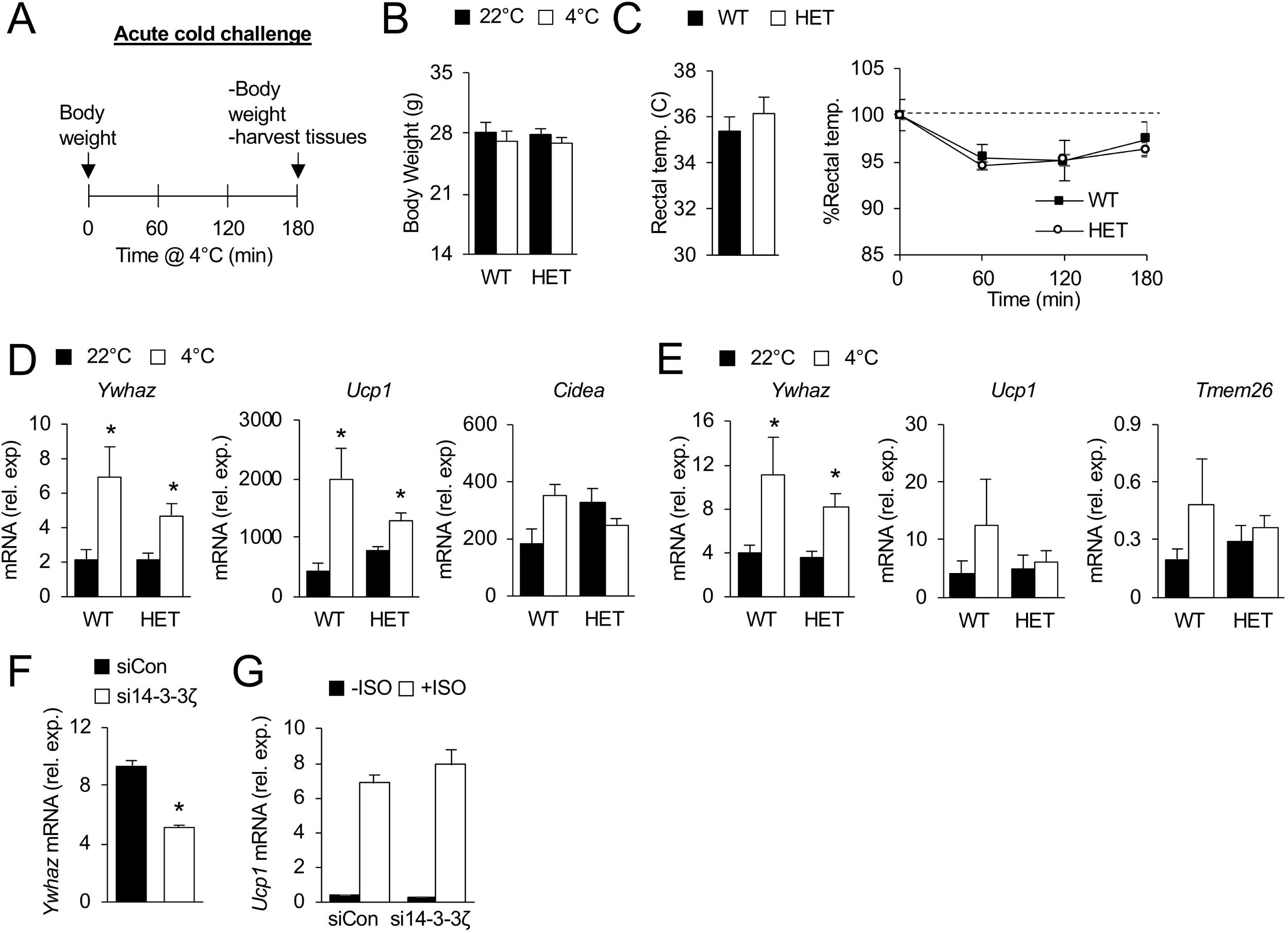
Loss of one allele of *Ywhaz* does not affect tolerance to acute cold. **(A)** Wildtype (WT) and mice lacking one allele of *Ywhaz*, the gene encoding 14-3-3ζ (HET) were challenged with cold for 3 hours. Temperature was measured by rectal probes at each time point. **(B, C)** Body weights (B) and rectal temperatures (C) of WT and HET mice were obtained prior, during, and at the end of the 3 hours cold challenge (n=7 mice per group). **(D**,**E)** Expression of brown-selective (D) and beige-selective (E) genes from BAT and iWAT, respectively, at room temperature (22 °C) and after 3 hours cold exposure (n=7 per group, *: p<0.05 when compared to 22 °C). **(F**,**G)** Knock-down of 14-3-3ζ expression by siRNA (F) in UCP1-luciferase (UCP1-Luc) cells, a brown adipocyte cell line, does not affected isoproterenol (ISO, 10μM, 4 hours)-mediated induction of *Ucp1* expression (G) (n=6 per group, *: p<0.05 when compared to -ISO; #: p<0.05 when compared to siCon). Data are represented as mean ± SEM.

To further determine if 14-3-3ζ is necessary for *Ucp1* mRNA expression in brown adipocytes, we utilized the UCP1-luciferase adipocyte cell line, an *in vitro* model of brown adipocytes (Galmozzi et al., 2014). Transient knockdown of *Ywhaz* by siRNA did not affect isoproterenol-induced *Ucp1* mRNA expression, which suggested that 14-3-3ζ is not required for *Ucp1* expression (Figure 1F,G). When taken together, these *in vivo* and *in vivo* data demonstrate that reducing 14-3-3ζ expression does not affect cold tolerance or *Ucp1* gene expression.

### 14-3-3ζ over-expression provides tolerance to acute cold

We next examined if increasing 14-3-3ζ expression could affect cold tolerance. 12-week-old transgenic (TAP) mice over-expressing a TAP-tagged human 14-3-3ζ molecule were challenged with acute cold (4°C) exposure for 3 hours (Figure 2A). In male mice, no differences in body weight were detected before or after cold exposure (Figure 2B); however, male TAP mice displayed a significant, restoration of rectal temperature, signifying improved tolerance to acute cold exposure (Figure 2C). A similar increase in body temperature was observed in female mice, but it was not statistically significant (Figure S2B). In BAT, acute cold exposure had no effect on the expression of *Ucp1, Pgc1a*, or *Cidea* (Figure 2D). In contrast, significant increases in *Ucp1* mRNA levels in iWAT of both male and female TAP mice following acute cold were detected (Figure 2E, S2D). Taken together, these data demonstrate that 14-3-3ζ over-expression is sufficient to improve tolerance to acute cold exposure.

**Figure 2.**
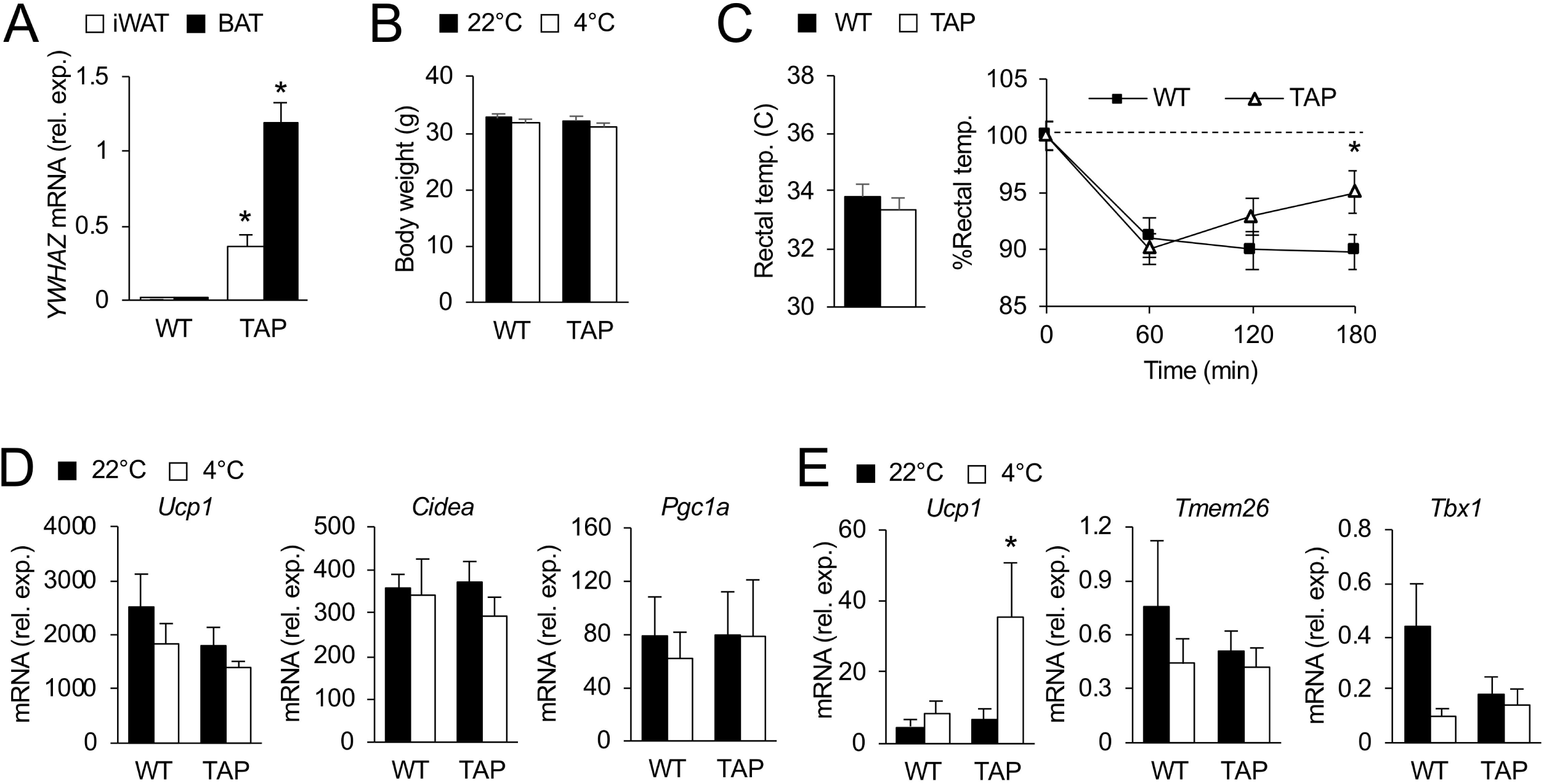
Over-expression of 14-3-3ζ enhances tolerance to acute cold in male mice. **(A)** Quantitative PCR was used to measure *YWHAZ* (gene encoding human 14-3-3ζ) in iWAT and BAT of TAP mice (n=3 per group, *: p<0.05). **(B**,**C)** Body weights (B) and rectal temperatures (C) of WT and TAP mice were obtained prior, during, and at the end of the 3 hour cold challenge (n=7 mice per group). **(D**,**E)** Expression of brown-selective (K) and beige-selective (L) genes from BAT and iWAT, respectively, at room temperature (22 °C) and after 3 hours cold exposure (n=7 per group, *: p<0.05 when compared to 22 °C). Data are represented as mean ± SEM.

### Over-expression of 14-3-3ζ improves tolerance to prolonged cold exposure

Given the improved tolerance to acute cold exposure in TAP mice and increased signs of beiging of iWAT (Figure 2C,E), male WT and TAP mice were subjected to a prolonged cold (4°C) challenge of 3 days (Figure 3A). Mice had similar body weights before cold exposure and lost comparable body weight after the challenge (Figure 3B). Moreover, fat and lean mass in WT and TAP mice before and after cold exposure were not different (Figure 3C). Additionally, no visible differences in the pelage, or fur, of mice were observed (data not shown). Food intake was increased in both WT and TAP mice when temperature was lowered from 22°C to 4°C, which is consistent with the need to supply necessary fuel as compensation for energy dissipated during thermogenesis (Bartelt et al., 2011), but no differences were observed between TAP and WT mice (Figure 3D). Moreover, locomotor activity or RER were not changed between groups (Figure 3E,F). With respect to core body temperature, TAP mice exhibited significantly higher body temperatures during the last 48 hours of cold exposure (Figure 3G,H), and unexpectedly, significantly decreased energy expenditure during the last 2 dark phases of the cold challenge were observed (Figure 3I,J).

**Figure 3.**
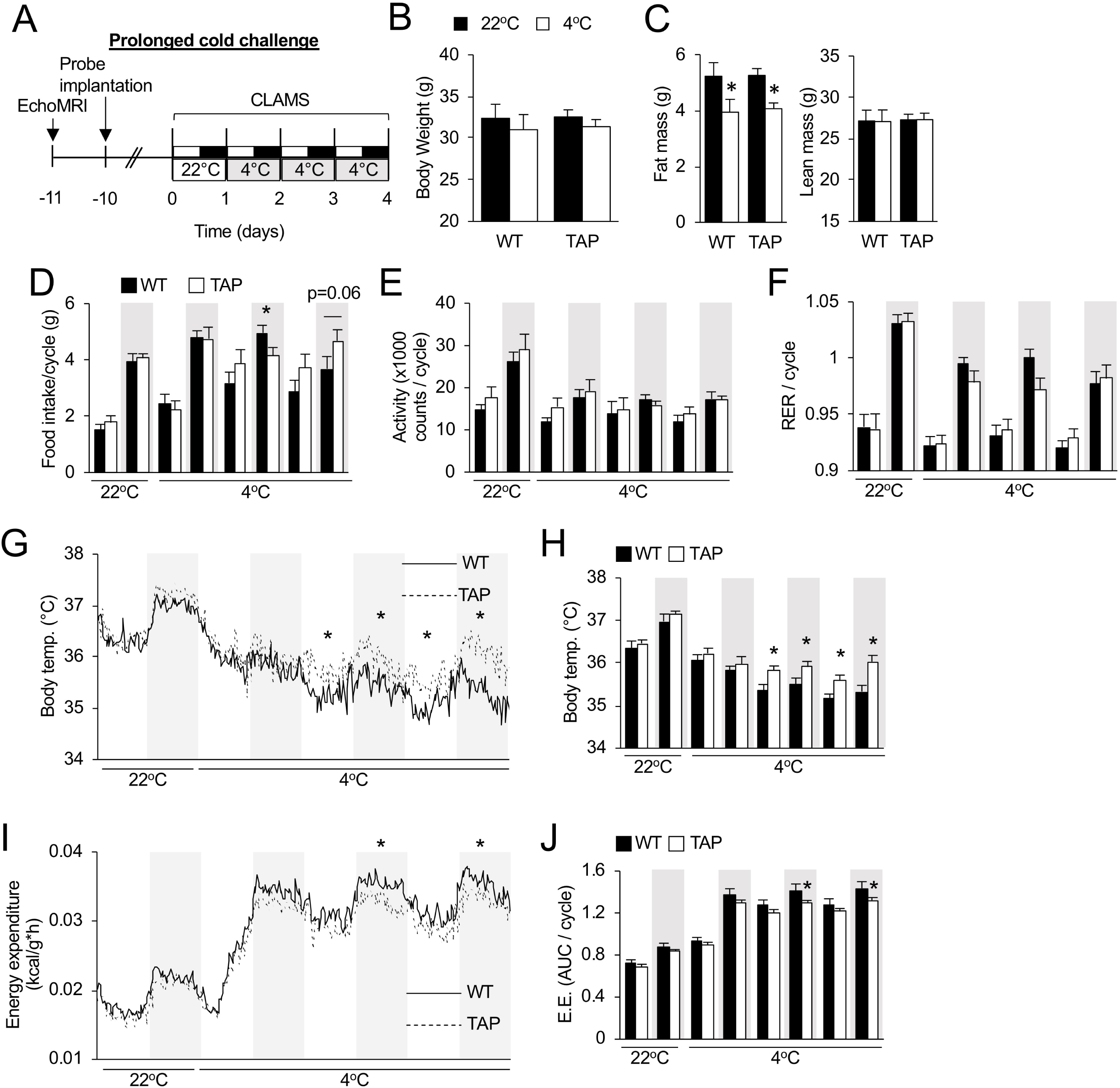
Over-expression of 14-3-3ζ improves tolerance to prolonged cold exposure in male mice. **(A)** Wildtype (WT) and 14-3-3ζ over-expressing (TAP) mice were implanted with a temperature probe 1 week prior to placement in CLAMS metabolic cages for 3 days at 4 °C. **(B**,**C)** Body weights (B) and lean and fat mass measurements by echo-MRI (C) were obtained prior to and after the prolonged cold challenge. **(D-F)** During the light and dark cycles at 22°C and 4 °C, food intake (G), locomotor activity (H), and RER were measured in WT and TAP mice. **(G-J)** Body temperature (G,H) and energy expenditure (I,J) were measured and are reported either as the average trace for all mice over the cold challenge (G, I) or as an average per light:dark cycle (H,J) (n=8 WT and n=10 TAP mice; *: p<0.05 when compared to WT). Data are represented as mean ± SEM.

Analysis of adipocyte morphometry revealed no differences in adipocyte size in iWAT or gonadal WAT (gWAT) of WT and TAP mice (Figure 4A-F). However, after prolonged cold exposure, distinct morphological differences were observed in BAT, whereby smaller lipid droplets were visible in TAP mice (Figure 4G). Despite these differences in the size of lipid droplets in BAT of TAP mice, circulating free fatty acids (FFAs) and glycerol levels following prolonged cold exposure were similar between WT and TAP mice (Figure 4H), and no differences in total triacylglycerol content in BAT were detected (Figure 4I).

**Figure 4.**
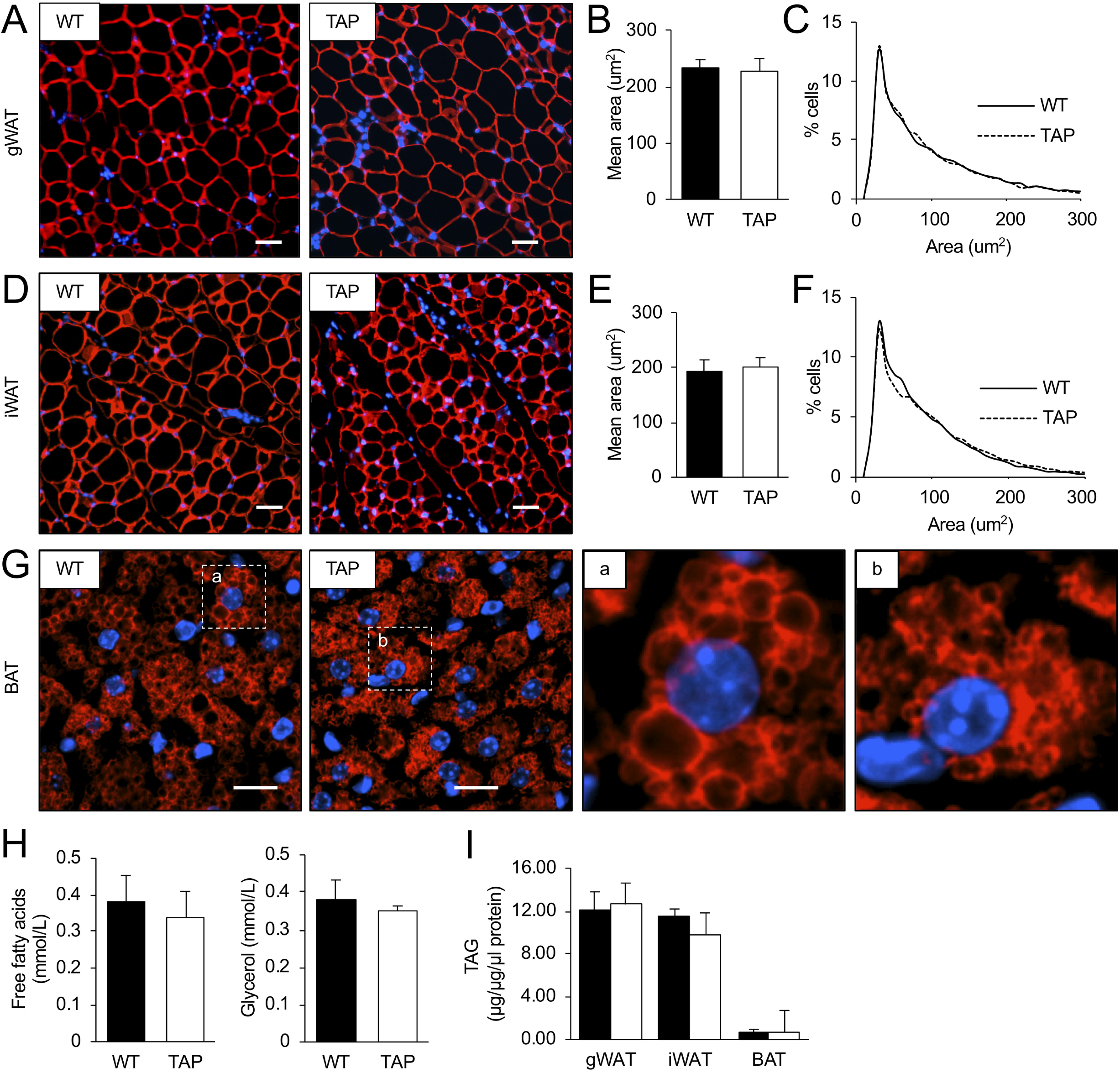
Adipocyte size in gonadal or inguinal white adipose tissue is not affected by 14-3-3ζ over-expression following cold exposure. **(A-F)** Immunofluorescent staining was used to examine adipocyte morphology (A,D), average size (B,E), and size distribution (C,F) in gonadal white adipose tissue (gWAT, A-C) and inguinal WAT (E,F) (Representative images of n=4-5 mice per group; scale bar= 200 μm,). (**G)** Morphology of BAT was measured by immunostaining for perilipin. Inset images of brown adipocytes from WT (a) and TAP (b) male mice (Representative images of n=4-5 mice per group; scale bars = 100 μm). **(H**,**I)** Circulating free fatty acids and triglycerides (H) and total triacylglycerols in gWAT, iWATm and BAT (I) of male WT and TAP mice were measured after 3 days of cold exposure (n= 4-5 per group). Data are represented as mean ± SEM.

### Over-expression of 14-3-3ζ increases beiging of inguinal white adipocytes

To better understand how 14-3-3ζ could improve tolerance to prolonged cold exposure, we first examined whether increased signs of beiging or BAT activity occurred as a result of 14-3-3ζ over-expression. In BAT, *Ucp1* mRNA was not different between groups, but differences in the expression of various thermogenic genes, such as *Prdm16 and Pdk4*, were detected between WT and TAP mice (Figure 5A). In iWAT of prolonged cold exposed animals, a five-fold increase in *Ucp1* mRNA levels was detected in TAP mice, in addition to significantly higher levels of *Tbx1* mRNA (Figure 5B). Marked elevations in UCP1 protein abundance were also detected in iWAT and BAT of TAP mice when compared to littermate controls (Figure 5C,D), and increased *Fgf21* mRNA levels could also be detected in cold-exposed iWAT from TAP mice (Figure 5E). Taken together, these findings indicate that 14-3-3ζ over-expression promotes the beiging of iWAT during prolonged cold exposure, which may account for the increased body temperature and improved cold tolerance.

**Figure 5.**
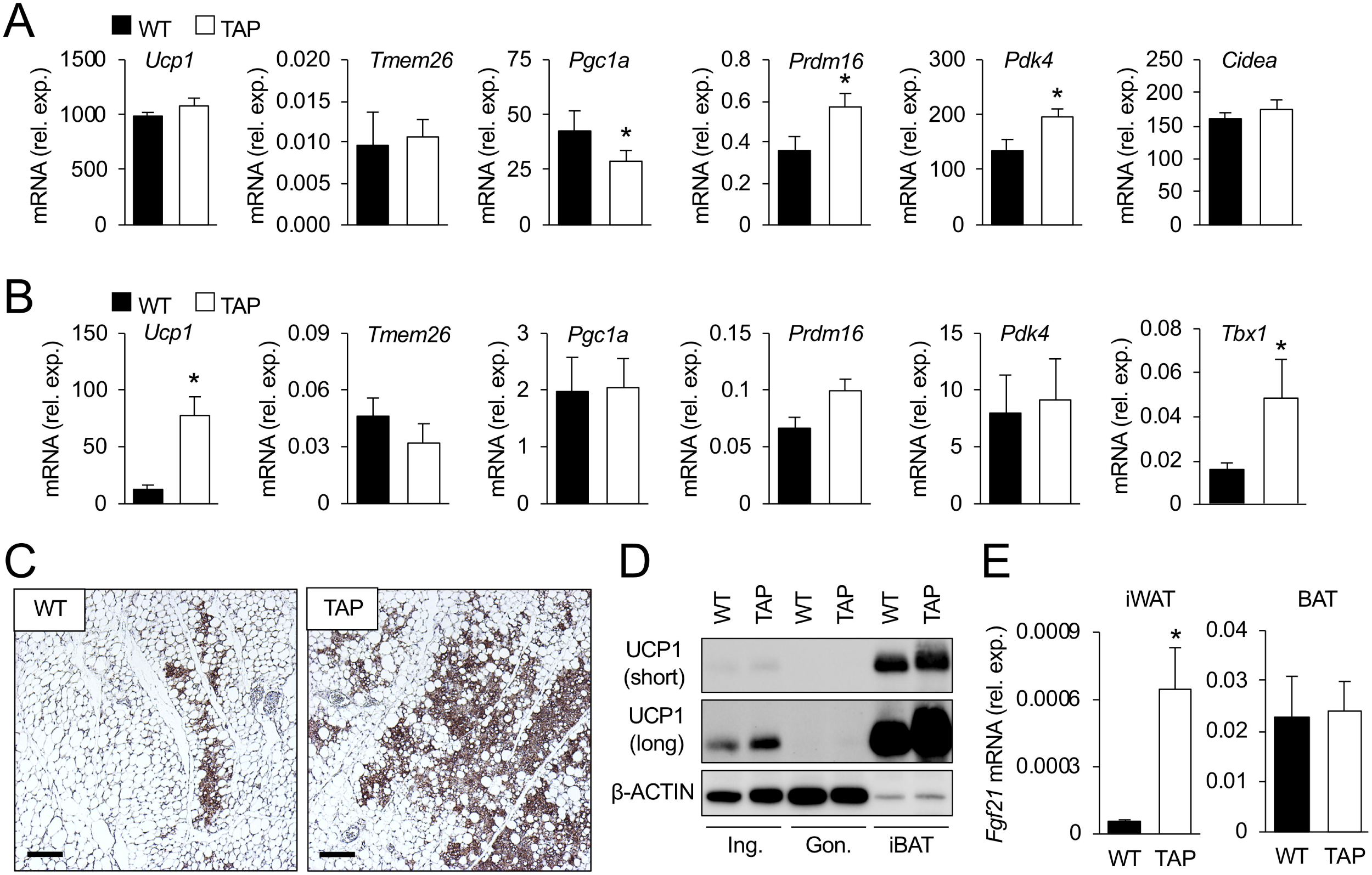
Transgenic over-expression of 14-3-3ζ increases beiging of ingulnal white adipocytes in male mice. **(A**,**B)** Following 3 days of cold exposure, brown adipose tissue (BAT, A) and inguinal white adipose tissue (iWAT, B) from wildtype (WT) and transgenic mice over-expressing 14-3-3ζ (TAP) were harvested, followed by quantitative PCR to measure genes associated with thermogenesis and beiging (n=8 WT and 10 TAP; *: p<0.05). **(C)** Immunohistochemistry was used to detect UCP1 immunoreactivity in paraffin embedded iWAT sections from WT or TAP mice exposed to prolonged cold (representative images for n=8 WT and 10 TAP mice; scale bar= 50 μm). **(D)** Immunoblotting was used to detect UCP1 protein in lysates from iWAT, gWAT, or BAT of prolonged cold-exposed WT and TAP mice (representative image of n=6 mice per genotype). **(E)** *Fgf21* mRNA levels were measured from iWAT harvested from WT and TAP mice exposed for 3 days to (n=6 per group, *: p<0.05). Data are represented as mean ± SEM.

### Over-expression of 14-3-3ζ does not alter sensitivity to β-adrenergic stimuli or sympathetic activity

During long-term cold exposure, the sympathetic nervous system releases norepinephrine, which acts as the principal activator of thermogenesis (Bartness et al., 2010; Cannon and Nedergaard, 2004; Fischer et al., 2017; Gabaldon et al., 2003; Landsberg et al., 1984; Thomas and Palmiter, 1997). Thus, to explore the possibility that altered sensitivity to β-adrenergic stimuli could account for differences in energy expenditure (Figure 3K,L), WT and TAP mice were chronically injected (7 days) with 0.9% saline or the β3-adrenergic receptor agonist CL-316,243 (CL, 1mg/kg) (Figure 6A). No differences in response to CL-mediated changes in total body weight, lean mass, or fat mass were observed between TAP and WT mice (Figure 6B,C). Levels of *Ucp1* mRNA were similarly increased by CL treatment in iWAT (Figure 6D) and BAT (Figure 6E) of both groups. Furthermore, markers of brown, *Cidea* and *Pdk4*, and beige, *Tmem26* and *Tbx1*, adipocytes were not different between TAP mice and WT littermate controls (Figure 6D,E). Taken together, these data suggest that over-expression of 14-3-3ζ does not alter sensitivity to β-adrenergic stimuli.

**Figure 6.**
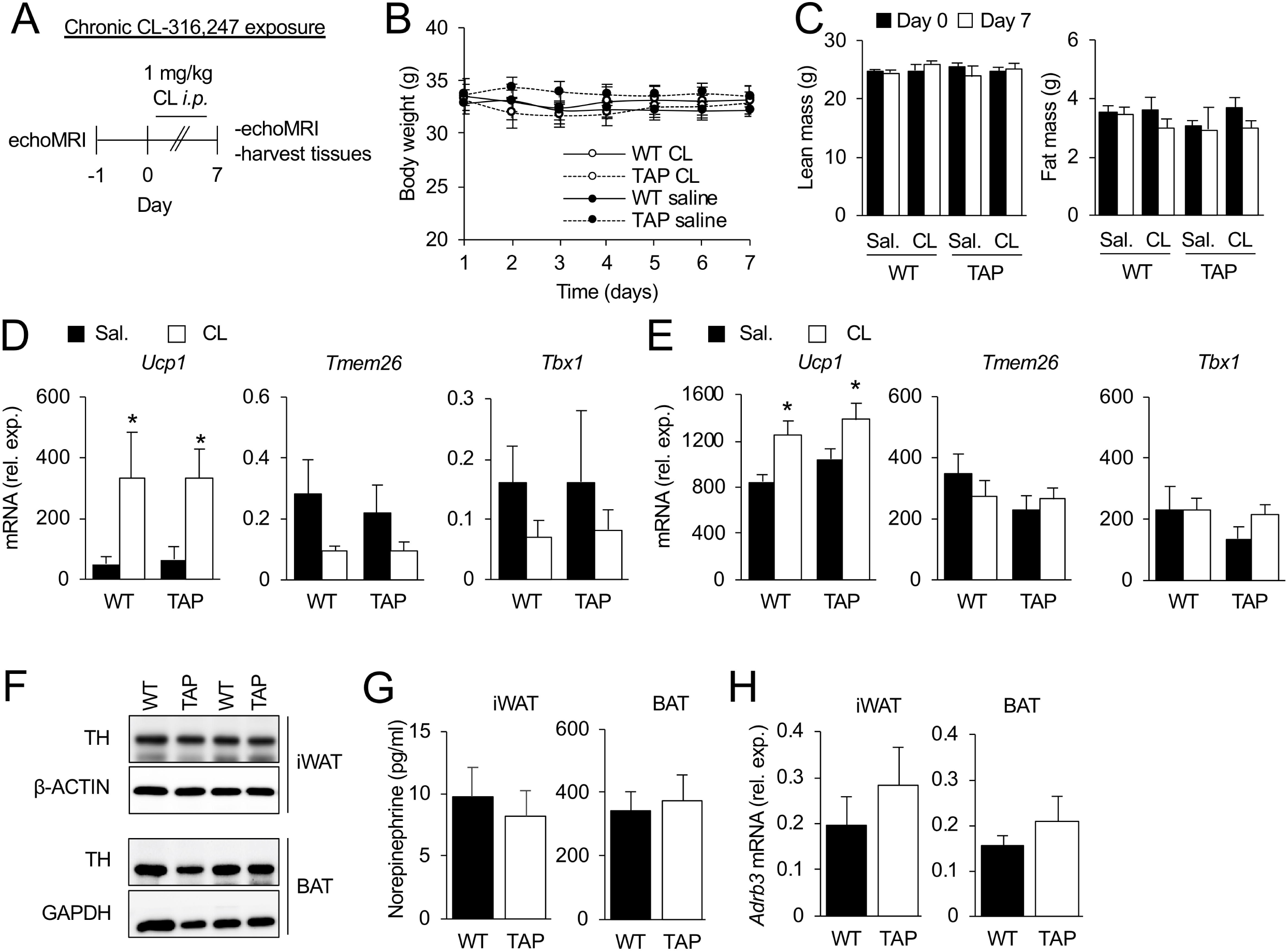
No differences in sympathetic activity or sensitivity were detected in mice over-expressing 14-3-3ζ. **(A)** To examine if 14-3-3ζ over-expression confers increased sensitivity to β-adrenergic stimuli, wildtype (WT) and transgenic 14-3-3ζ over-expressing mice (TAP) were injected with the β3-adrenergic agonist CL,316,247 (CL, 1 mg/kg) for seven days. Lean and fat mass were measured by echoMRI before and after CL injections. **(B**,**C)** Body weight (B) and lean and fat mass (C) were obtained before and after CL-treatment. **(D, E)** Inguinal white adipose tissue (iWAT, D) and brown adipose tissue (BAT, E) were harvested from WT and TAP mice after 7 days of injections with saline (sal.) or CL exposure to measure genes associated with thermogenesis. **(F-H)** Cold-exposed iWAT and BAT were harvested from WT and TAP mice to measure protein abundance of tyrosine hydroxylase by immunoblotting (F), norepinephrine by ELISA, and *Adrb3* mRNA levels by quantiative PCR (H). (n=5 WT and 7 TAP, *: p<0.05 when compared Day=0). Data are represented as mean ± SEM.

The above *in vivo* studies demonstrate that 14-3-3ζ-mediated cold adaptation is not due to increased sensitivity to adrenergic stimuli. Thus, we turned our focus to investigate whether changes in sympathetic innervation or activity in iWAT or BAT of TAP mice could be occurring (Wang et al., 2019). Tyrosine hydroxylase (TH) protein expression was not altered in iWAT or BAT (Figure 6F) of both groups following prolonged cold exposure, and consistent with these observations, norepinephrine levels in iWAT and BAT were not different between WT and TAP mice (Figure 6G), nor *were* mRNA levels of *Adrb3* expression in iWAT and BAT (Figure 6H). Together, these findings suggest that sympathetic innervation or activity is not altered in TAP mice in response to prolonged cold exposure.

### 14-3-3ζ does not influence glucose utilization or oxidative metabolism during prolonged cold exposure

Brown and beige adipocytes utilize triglycerides and glucose as sources of energy for heat production (Kajimura et al., 2015; Wang and Seale, 2016), and given the differences in cold tolerance between WT and TAP mice, we examined whether glucose handling was altered by systemic 14-3-3ζ over-expression. As a first step, we examined whether differences in glucose tolerance could be detected in WT and TAP mice following three day exposure of mice to 4°C cold. Following a 6-hour fast, no differences in fasting body weights or blood glucose levels were observed between WT and TAP mice (Figure 7A,B), and at 22°C, TAP mice had improved glucose tolerance when compared to WT mice, but this improvement was lost following prolonged cold exposure (Figure 7C).

**Figure 7:**
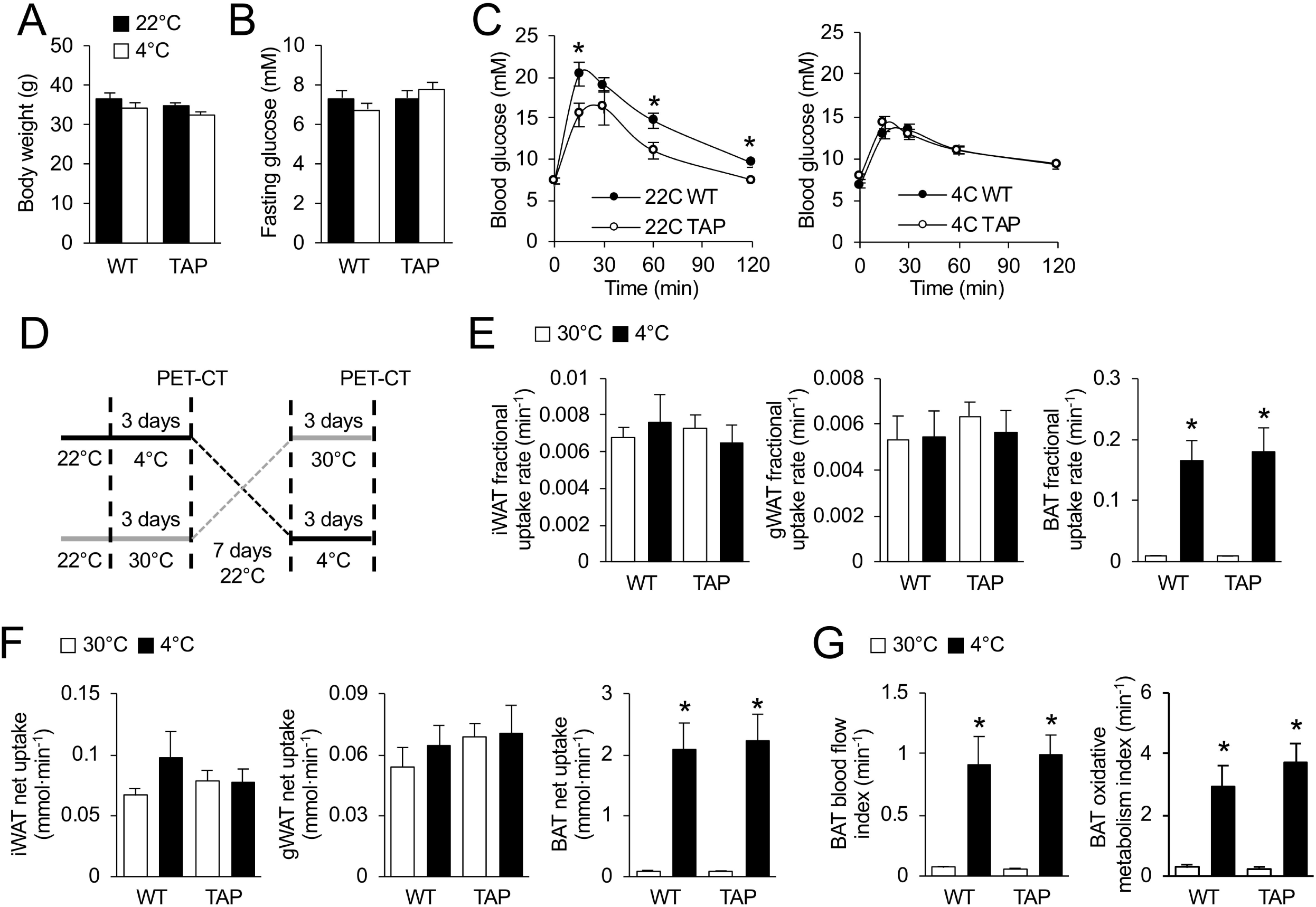
Over-expression of 14-3-3ζ does not enhance the metabolic activity of BAT during cold exposure. **(A-C)** Wildtype (WT) or transgenic mice over-expressing 14-3-3ζ (TAP) housed at 22°C or 4°C did not display differences in body weight (A) or fasting blood glucose (B) prior to an intraperitoneal glucose tolerance test (C, 2 g/kg) (n=8 WT and 6 TAP; *: p<0.05). **(D)** To measure the metabolic activity of iWAT and BAT during prolonged cold exposure, WT and TAP mice were subjected to a randomized cross-over design whereby mice were housed at thermoneutrality (30°C) or 4°C for 3 days, with 1-week recovery in between temperatures, and subjected to dynamic [^18^F]-FDG and [^11^C]-acetate-based PET-CT imaging. **(E**,**F)** Fractional and net [^18^F]-FDG uptake was measured in BAT, gWAT, and iWAT from WT and TAP mice at the indicated temperatures (n=8 WT, 10 TAP; *: p<0.05). **(G)** Oxidative activity, as measured by [^11^C]-acetate metabolism, and blood flow were measured in BAT of WT and TAP mice at the indicated temperatures (n=8 WT, 10 TAP; *: p<0.05). Data are represented as mean ± SEM.

To better understand whether 14-3-3ζ over-expression altered glucose utilization or metabolic activity in BAT or iWAT, [^18^F]-FDG and [^11^C]-acetate-based PET-CT imaging was used. Mice were subjected to a randomized cross-over study where they were housed at thermoneutrality (30°C) or 4°C for 3 days with a week recovery in between temperatures, followed by dynamic PET-CT imaging (Figure 7D). Analysis of fractional and net [^18^F]-FDG uptake in non-adipose tissues revealed no differences between groups (Figure S3). Of note was the observation that [^18^F]-FDG uptake in muscle was elevated in WT mice at 4C, but not in TAP mice (Figure S3B). When considering the glucose excursion profiles on WT and TAP mice following an *i*.*p*. glucose bolus (Figure 7C), these findings demonstrate that TAP mice do not depend on enhanced glucose uptake as an energy source in response to prolonged cold. Furthermore, no differences in fractional and net [^18^F]-FDG uptake were observed in BAT, iWAT, or gWAT (Figure 7E,F).

With respect to oxidative metabolism rates among non-adipose tissues, only myocardial tissue displayed decreased metabolic activity (Figure S4). Oxidative metabolism increased in BAT of WT and TAP mice following cold exposure; however, no significant differences were observed between groups (Figure 7G). Oxidative metabolic rates in iWAT were too low for detection (data not shown). When taken together, the absence of differences in gucose uptake or metabolic activity of BAT or iWAT suggest an alternative mechanism must be contributing to the increased body temperature observed in TAP mice during prolonged cold exposure.

### 14-3-3ζ increases vasoconstriction to mitigate heat loss

The observed increases in body temperature and paradoxical decreases in energy expenditure suggest that an alternative, adaptive mechanism of heat conservation must be present in TAP transgenic mice. To further explore this phenomenon, thermal conductance, a measurement of the rate of heat dissipation to the environment (Kaiyala et al., 2016), was determined in WT and TAP at 22°C and 4°C. In contrast to WT mice, TAP mice displayed significantly lower thermal conductance at room temperature and throughout the prolonged cold exposure period (Figure 8A). This suggests that an alternative, separate mechanism is active in TAP mice to mitigate heat loss during mild (22°C) and severe cold (4°C) stress. Leptin was previously found to have thermo-regulatory effects in *ob/ob* mice; however, circulating levels of leptin were not different between WT and TAP mice (Figure 8B) (Fischer et al., 2016; Kaiyala et al., 2016; Kaiyala et al., 2015). To explore whether 14-3-3ζ over-expression affected vasoconstriction *in vivo*, laser doppler imaging was used to measure superficial blood-flow as a surrogate measure of vessel diameter (Thomas et al., 1994). At 22°C, blood flow was significantly reduced in TAP animals (Figure 8C), which suggests increase vasoconstriction. Taken together, these findings suggest that 14-3-3ζ can activate alternative mechanisms to conserve heat loss and maintain homeothermy.

**Figure 8:**
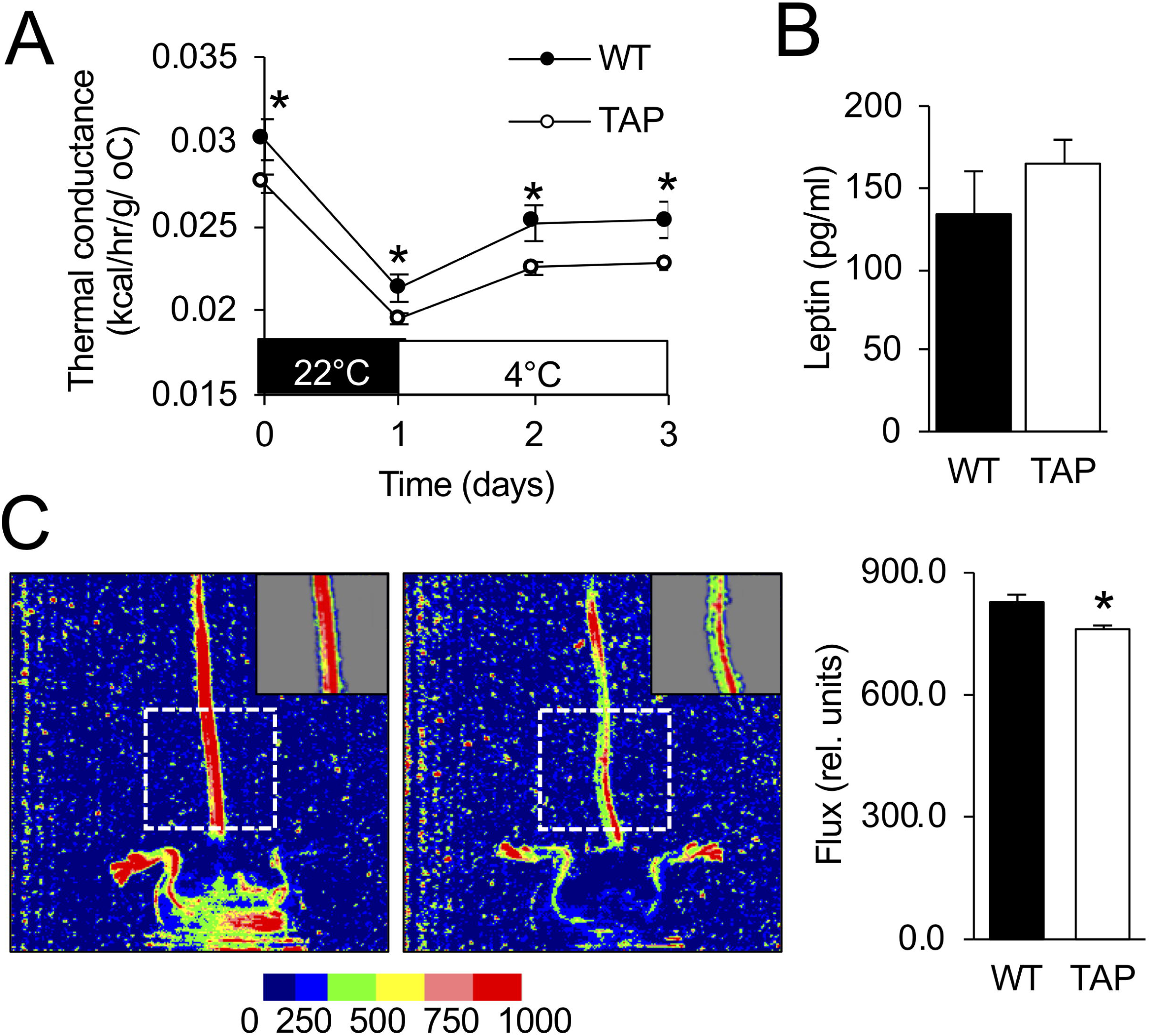
Increased heat retention is associated with 14-3-3ζ over-expression. **(A)** Thermal conductance was calculated for WT and TAP mice housed at 22°C or 4°C during the prolonged cold exposure CLAMS study (Figure 2) (n=8 WT and 10 TAP mice; *: p<0.05 when compared to WT). **(B)** Plasma leptin was measured from blood samples obtained from WT and TAP mice after the prolonged cold challenge (n=8 WT and 10 TAP mice). **(C)** Laser doppler imaging was used to measure blood flow in the base of the tail of WT and TAP mice housed at 22°C. (n=16 WT and 22 TAP mice; *: p<0.05). Data are represented as mean ± SEM.

## Discussion

Substantial interest in understanding the roles of beige and brown adipocytes in the regulation of homeothermy and energy homeostasis has occurred over the past decade, and due to their abilities to metabolize lipids via β-oxidation, this has sparked interest in exploring the possibility of activating beige and brown adipocytes as a therapeutic approach to treat obesity and diabetes. In the context of homeothermy, most studies assume UCP-1-dependent thermogenic pathways to be the priniciple mechanism to protect against cold, and little emphasis is placed on alternative, UCP-1-independent mechanisms to influence cold tolerance. Herein, we demonstrate that over-expression of 14-3-3ζ is sufficient to improve tolerance to both acute and prolonged cold exposure. Despite the ability of mice over-expressing 14-3-3ζ to raise body temperature to defend against cold, this physiological response is associated with a parallel decreases in energy expenditure and the restriction of peripheral blood-flow to retain heat. By minimizing heat loss, less energy has to be expended to defend against hypothermia and maintain body temperature. It should be noted that 14-3-3ζ over-expression was not associated with changes in sympathetic innervation or sensitivity. Taken together, these findings demonstrate that while thermogenic mechanisms may occur by way of increased BAT activity or the beiging of white adipocytes, alternative mechanisms that promote heat conservation during cold adaptation must also be considered in the context of cold tolerance.

Adaptive mechanisms to maintain core body temperature are activated to defend against cold exposure, and this predominantly results in the production of heat via UCP1-dependent mechanisms in brown and beige adipocytes (Cannon and Nedergaard, 2004; Morrison and Nakamura, 2019). However, in the present study, the improved tolerance of transgenic mice over-expressing 14-3-3ζ to prolonged cold was associated with decreased energy expenditure, despite the ability of transgenic mice to raise and maintain higher body temperatures. Intact adaptive thermogenic mechanisms are present in TAP mice, as the expected increases in UCP1 expression in BAT and beige iWAT following cold exposure cold be detected, as well as the increase in oxidative metabolism of BAT. Thus, an alternative mechanism to defend against changes in core body temperature that are independent of thermogenesis must be active in TAP mice over-expressing 14-3-3ζ.

Thermogenesis is largely viewed as the primary mechanism for cold tolerance, but when this process is disrupted, as in the case with UCP1-deficient mice, vasoconstriction becomes the predominant mechanism for cold adaptation (Wang et al., 2006). The decrease in thermal conductance during mild (22°C) and severe (4°C) cold stress raised the possibility that 14-3-3ζ over-expression could activate processes, such as vasoconstriction, to mitigate heat loss from skin (Kaiyala et al., 2016). Indeed, we found that 14-3-3ζ over-expression was associated with reduced peripheral blood-flow in tails of TAP mice, which is indicative of vasoconstriction (Wang et al., 2006). Leptin has been recently identified as having vasoconstrictive effects, but no differences in circulating leptin levels were detected between WT and TAP mice (Fischer et al., 2016; Kaiyala et al., 2016; Kaiyala et al., 2015). It is possible that the increase in vasoconstriction could be due to elevated levels of norepinephrine, a potent vasoconstrictor, in the circulation, but it is also possible that 14-3-3ζ over-expression could have direct effects in endothelial cells. For example, 14-3-3ζ has been found to interact with α2A-adrenergic receptors subtypes at the i3 loop, which blocks their interactions with β-arrestins to limit their internalization. Increased expression at the cell membrane of endothelial cells could result in potentiated α2A-adrenergic receptor-mediated effects on vasoconstriction (Prezeau et al., 1999; Wang and Limbird, 2002). Thus, 14-3-3ζ over-expression could havev effects on receptor density at the cell membrane of endothelial cells.

As over-expression of 14-3-3ζ was not associated with changes in food intake or locomotor activity during cold exposure, the observed decrease in energy expenditure could eventually lead to a long-term imbalance in energy homeostasis, thus increasing the propensity to development of obesity (Bluher, 2019; Rodgers et al., 2012). Indeed, we previously demonstrated that transgenic over-expression of 14-3-3ζ was sufficient to exacerbate age- and high-fat diet-induced weight gain (Lim et al., 2015). Moreover, elevated expression of 14-3-3ζ has been detected in visceral adipose tissue from individuals with obesity (Capobianco et al., 2012; Insenser et al., 2012). Whether the increase in adiposity due to elevations in 14-3-3ζ expression is due to changes in overall metabolism or cell-autonomous effects in adipocyte development require further in-depth investigation, and it is necessary to examine whether increasing 14-3-3ζ expression could have other effects on age-associated declines in metabolism and overall health (Mills et al., 2016).

In summary, we demonstrate in this study that over-expression of 14-3-3ζ in male mice is sufficient to increase tolerance to cold. Despite the activation of thermogenic mechanisms, we demonstrate that cold-induced increases in body temperature are paradoxically associated with decreased energy expenditure in mice over-expressing 14-3-3ζ and increased peripheral vasoconstriction to retain heat. This suggests that alternative, non-thermogenic mechanisms can also be activated to defend against cold and raise the notion that other mechanisms are equally important in the defence of homeothermy and should be considered when studying cold tolerance. Taken together, our results show an important role of 14-3-3ζ in adaptation to cold and add new insights to the regulation of key physiological processes by molecular scaffolds.

## Methods

### Animal studies

Wildtype (WT) and 14-3-3ζ heterozygous (HET) mice were on a C57Bl/6J background, whereas transgenic (TAP) mice over-expressing a TAP-tagged human 14-3-3ζ molecule were on a CD-1 background (Angrand et al., 2006; Lim et al., 2015). Mice were housed in a temperature-controlled (22°C) animal facility on a 12-hour light/dark cycle in the Centre de Recherche du Centre Hospitalier de l’Université de Montréal (CRCHUM). Mice had *ad libitum* access to water and standard chow (TD2918, Envigo, Huntingdon, United Kingdom), and all animal studies were performed in accordance to the Comité Institutionnel de Protection des Animaux (CIPA) of the CRCHUM.

For acute cold challenges, 12 week-old mice were individually caged and fasted for 4 hours prior to and during a 3-hour challenge at 4°C with *ad libitum* access to water. Body temperature was measured with a physio-suit rectal probe (Kent scientific, Torrington, CT, USA). For prolonged cold challenges, mice were housed in Comprehensive Lab Animal Monitoring System (CLAMS, Columbus Instruments Columbus, OH, USA) metabolic cages or a cold room for 3 days at 4°C. The β3 adrenergic agonist CL316,243 (Sigma Aldrich, St Louis, MO, USA) was diluted in saline 0.9%, and mice received daily intraperitoneal injections of either saline 0.9% or CL316,243 (1mg/kg) for 7 days. Body composition (lean and fat mass) was determined using EchoMRI (EchoMRI™, Houston, TX, USA).

### Cell culture

The immortalized UCP1-luciferase (UCP1-Luc) adipocyte cell line was kindly provided by Dr. Shingo Kajimura (Diabetes Center, University of California-San Francisco) (Galmozzi et al., 2014). Cells were grown in 25 mM glucose DMEM media (Thermo Fisher Scientific, Waltham, MA, USA), supplemented with 10% fetal bovine serum FBS (Thermo Fisher Scientific) and 1% streptomycin (Thermo Fisher Scientific), and grown at 37°C, 5% CO_2_. UCP1-Luc cells were induced to differentiation into brown adipocytes with a cocktail containing 5μg/ml insulin (Sigma Aldrich), 0.5mM 3-Isobutyl-1-methylxanthine IBMX (Sigma Aldrich), 1μM dexamethasone (Sigma Aldrich), 0.125mM indomethacin (Sigma Aldrich), and 1nM 3,3′,5-Triiodo-L-thyronine T3 (Sigma Aldrich) for 2 days, followed by maintenance medium containing 5μg/ml insulin and 1nM T3 on day 3, and DMEM/FBS growth medium on day 5. Lipofectamine RNAiMax (Invitrogen, Carlsbad, CA,USA), and Silencer Select siRNA against *Ywhaz* (gene encoding 14-3-3ζ) and a scrambled, control siRNA (Ambion, Austin, TX, USA) were used to knockdown 14-3-3ζ, as previously described (Lim et al., 2015).

### Immunoblotting

Inguinal (iWAT) and gonadal (gWAT) white adipose tissues and interscapular brown adipose tissue (BAT) were homogenized in RIPA lysis buffer (50 mM β glycerol phosphate, 10mM Hepes, pH=7.4, 70 mM NaCl, 1% Triton X-100, 2mM EGTA, 1mM Na_3_VO_4_, 1mM NaF), supplemented with protease and phosphatase inhibitors. Lysates were centrifuged at 13000 rpm for 15 minutes at 4°C, the supernatant was collected, and protein concentration was determined using Bio-Rad protein assay dye Reagent (Bio-Rad, Hercules, CA, USA). Protein samples were resolved by SDS-PAGE, transferred to PVDF membranes and blocked with I-block (Applied Bio-systems, Foster city, CA, USA) for 1 hour at room temperature, followed by overnight incubation at 4 °C with primary antibodies against UCP1 (1:1000, R&D systems, Minneapolis, MN, USA), 14-3-3ζ (1:1000 Cell Signaling, Danvers, MA, USA), β-Actin (1:10000, Cell Signaling), β-Tubulin (1:1000, Cell Signaling) and Tyrosine hydroxylase (1:1000, Millipore, Bilerica, MA, USA). On the next day, membranes were washed and incubated with horseradish peroxidase-conjugated secondary antibodies (1:5000, Cell Signaling) for 1 hour at room temperature. Immunoreactivity was detected by chemiluminescence with a ChemiDoc system (Bio-Rad). Information for each antibody can be seen in Supplemental Table 1.

### Histology and Immunofluorescence

IWAT, gWAT, and BAT were excised and fixed in 4% PFA (Sigma Aldrich) for 7 days and stored in 70% ethanol prior to embedding in paraffin. Sections at 5 μm thickness were deparaffinized, re-hydrated and stained with Hematoxylin (Sigma Aldrich) and Eosin (Sigma Aldrich). Alternatively, slides were stained with a UCP1 antibody (1:250, Abcam, Cambridge, United Kingdom), followed by a HRP-conjugated secondary antibody for DAB labeling (Cell signaling). Images were taken at 20X (Nikon Eclipse Ti2, Nikon Instruments Inc, Melville, NY, USA).

For immunofluorescence, sections were stained for Perilipin (1:400, Cell Signaling). Antigen retrieval was performed with 10 mM Sodium Citrate buffer (Sigma Aldrich) at pH=6-6.2 for 15 min at 95°C. Sections were blocked 1 hour at room temperature with PBS-T (0.1% Triton, 5% normal donkey serum) and incubated overnight in PBS-T at 4°C with primary antibodies. Alexa Fluor 594-conjugated secondary antibodies (Jackson Immuno-research laboratories, Inc, West grove, PA, USA) were incubated for 1 hour at room temperature, and slides were mounted in Vectashield containing DAPI (Vector laboratories, Burlingame, CA, USA). Total adipocyte number and area was counted from 8-10 images per mouse, then measured using the Cell Profiler software (CellProfiler Analyst, Stable (2.2.1) (Jones et al., 2008). All immunofluorescence pictures were acquired with an EVOS FL microscope (Thermo Fisher Scientific).

### RNA isolation and quantitative PCR

Total RNA was isolated from cells and tissues using the RNeasy Mini kit or the RNeasy Plus Mini Kit (Qiagen, Montreal, Quebec, Canada respectively, and stored at −80°C. Reverse transcription was performed with the High Capacity cDNA Reverse Transcription Kit (Applied Biosystems) or the Superscript VILO Kit (Invitrogen), in accordance with manufacturer’s instructions. Gene expression was analysed by quantitative PCR (qPCR) with SYBR Green chemistry (PowerUp SYBR, ThermoFisher Scientific) on a QuantStudio 6 Real Time PCR machine (Applied Bio Systems, Life Technologies, Carlsbad, CA, USA). Relative gene expression was normalized to the reference gene, *Hprt*, as previously described (Lim et al., 2015; Mugabo et al., 2018). For a complete list of primers and their respective sequences, please see Supplemental Table 2.

### Metabolic phenotyping

Male WT and TAP mice at 16 weeks of age received abdominal surgery to implant a temperature probe 10 days prior to their placement in CLAMS. Body weight and body composition were measured before and after prolonged cold exposure using EchoMRI on living, non-anesthetized mice. Mice were singly housed in CLAMS cages with *ad libitum* access to water and normal chow diet and were maintained on a 12-hour light/dark cycle on the following schedule: 24 hours at 22°C for acclimatization, 24 hours at 22°C for basal measurements, and 72 hours at 4°C for the prolonged cold challenge. Food intake, respiratory exchange ratio, locomotor activity (beam breaks), energy expenditure (heat), core body temperature (°C) were measured in real-time every 15-20 mins. Following prolonged cold exposure, blood and tissues were collected and snap frozen for subsequent use. Thermal conductance, or ease by which heat escapes to the environment was calculated, as previously described (Kaiyala et al., 2015). It is derived from the formula, C = EE/(Tb − Ta), where C = conductance; EE = energy expenditure; Tb = core temperature and Ta = ambient temperature (Kaiyala et al., 2015).

Cohorts of mice were administered intraperitoneal glucose tolerance tests (IPGTTs) after 6 hours of fasting. IPGTTs were performed at 22°C and after 3 days of 4°C exposure. Mice were injected with glucose (2 mg/kg), and blood glucose was measured with a Contour Next glucose meter (Ascensia Diabetes Care, Basel, Switzerland). Circulating free fatty acids were measured from plasma samples using the Wako NEFA-HR (2) assay kit (Wako Pure chemical Industries LTD, Osaka, Japan) and circulating glycerol was measured from plasma samples using the triglyceride and free glycerol reagents (Sigma Aldrich) as per to manufacturers’ instructions. Circulating leptin (ALPCO, Salem, NH, USA) was measured from plasma samples, in accordance to manufacturers’ protocols. Norepinephrine (Rocky Mountain Diagnostics, Colorado Springs, CO, USA) was measured from iWAT and BAT tissue extracts following manufacturers’ instructions.

### Laser doppler imaging

Laser doppler imaging was used to measure peripheral tail blood flow, as a surrogate for vasoconstriction (Thomas et al., 1994). In brief, female WT and TAP mice were anesthetized and tail perfusion was measured with a Laser Doppler Perfusion Imager (LDPI) System (Moor Instruments Ltd., Axminister, UK) (Haddad et al., 2009; Turgeon et al., 2012). Consecutive measurements were obtained from anesthetized mice by scanning the base of the tail. Color images were acquired, and flux values were measured by calculating the average perfusion signal. All experiments were performed at 22°C, and anesthetized mice were placed on heating pads until laser doppler measurements. Mice were sacrificed at the end of the study by exsanguination under anesthesia (isoflurane).

### Positron emission tomography/computed tomography (PET/CT) imaging

μPET/CT experiments were approved by the Ethical Committee for Animal Care and Experimentation of the Université de Sherbrooke. Mice underwent a randomized, cross-over thermoneutrality (30°C) *vs*. cold exposure (4°C) for 3 days, with a one-week washout period in between, prior to sequential μPET dynamic imaging with [^11^C]-acetate and [^18^F]-FDG (2-[^18^F]fluoro-2-deoxy-glucose) after 6-hr fasting. μPET/CT experiments were performed under anesthesia (isoflurane 2.0%, 1.5 L.min^-1^), delivered to the animal through a nose cone. After cold exposure, mice were injected with the β3-agonist CL316,243 (2 mg/kg) prior to the PET tracer injection to maintain cold-induced BAT stimulation, as previously published (Labbe et al., 2015). All PET tracers were injected through the tail vein, and imaging was performed with the avalanche photodiode-based small animal μPET scanner (LabPET/Triumph, Gamma Medica, Northridge, CA) in the Sherbrooke Molecular Imaging Centre (*Centre de recherche du Centre hospitalier universitaire de Sherbrooke*, Sherbrooke, QC, Canada). Anaesthetized mice were placed on the scanner bed and positioned with the heart centered within the field-of-view of the scanner. A bolus of [^11^C]-acetate (10 MBq, in 0.2 mL of 0.9% NaCl) was injected intravenously followed by a 15-min whole body dynamic μPET acquisition. Then, a bolus of [^18^F]-FDG (10 MBq, in 0.1 mL of 0.9% NaCl) was injected and a 20-min whole body dynamic PET acquisition was performed. Residual [^11^C]-acetate activity during [^18^F]-FDG acquisition was corrected by acquiring a 60 s frame prior to the injection of [^18^F]-FDG, accounting for the disintegration rate of [^11^C]. Low dose CT scan imaging was performed using the integrated X-O small-animal CT scanner of the Triumph platform, consisting of a 40 W X-ray tube with a 75 μm focal spot diameter and a 2240 × 2368 CsI flat panel X-ray detector. All images were analyzed as described previously (Labbe et al., 2015). Tissue (except BAT) blood flow index [K_1_ in min^-1^ of [^11^C]-acetate and oxidative metabolism index (K_2_ in min^-1^ of [^11^C]-acetate) were estimate from [^11^C]-acetate using a three-compartment model (Klein et al., 2001). BAT K_1_ and K_2_ were estimate from [^11^C]-acetate using a new multi-compartment model specific to BAT (Richard et al., 2019). Tissue glucose fractional extraction (Ki – *i*.*e*., the fraction of circulating glucose taken up by tissues over time) was determined using the Patlak graphical analysis of [^18^F] activity. Tissue glucose uptake (Km) was determined by multiplying Ki by the plasma glucose concentration.

### Statistical analysis

Data are presented as mean ± standard error. Statistical analyses were performed with GraphPad Prism 8 (GraphPad Software) using Student’s t-test or one- or two-way ANOVA, followed by appropriate *post-hoc* tests. Statistical significance was achieved when p < 0.05.

## Acknowledgements

The authors would like to thank Dr. Amparo Acker-Palmer (University of Frankfurt, Germany) for providing original breeding pairs of TAP mice. The authors would like to acknowledge the CRCHUM rodent metabolic phenotyping core facility for their help with CLAMS and echoMRI studies and the Molecular Pathology core histological support. The authors also thank Mélanie Archambault, Jean-François Beaudoin and Maxime Paillé (from the Sherbrooke Molecular Imaging Center, Sherbrooke, Quebec, Canada) for their technical assistance with µPET. The authors are grateful to Dr Ouhida Benrezzak for her assistance with all protocols related to animal care and ethics committee.Funding for this study was provided by a Canadian Institute of Health Research (CIHR) Project Grant (PJT-153144) to GEL, a Heart and Stroke Foundation of Canada grant-in-aid (HSFC G-17-0019106) to AR, CIHR grants to AC (MOP 53094 and 299962), and an Intercenter Structuring Initiative by the Réseau de recherche en santé CardioMétabolique, Diabète et Obésité (CMDO) to GEL. GEL is the Canada Research Chair in Adipocyte Development and AC is the Canada Research Chair in Molecular Imaging of Diabetes. KD was supported by a Stage Intercentre Grant by the CMDO and an International tuition fee waiver in virtue of a bilateral agreement between the government of Senegal and the government of Quebec.

## Contributions

K.D. designed the studies, carried out the research, interpreted the results, and wrote the manuscript. A.F.L. provided 14-3-3ζ Het mice. S.D. and A.R designed and carried out laser doppler analysis of blood flow and revised the manuscript. C.N and A.C.C. designed experiments, analyzed data, and wrote the manuscript. G.E.L conceived the concept, designed the studies, interpreted the results, and wrote and revised the manuscript. G.E.L is the guarantor of this work.

