## Supplemental Figures and Tables for "14-3-3ζ mediates an alternative, non-thermogenic mechanism to reduce heat loss and improve cold tolerance"

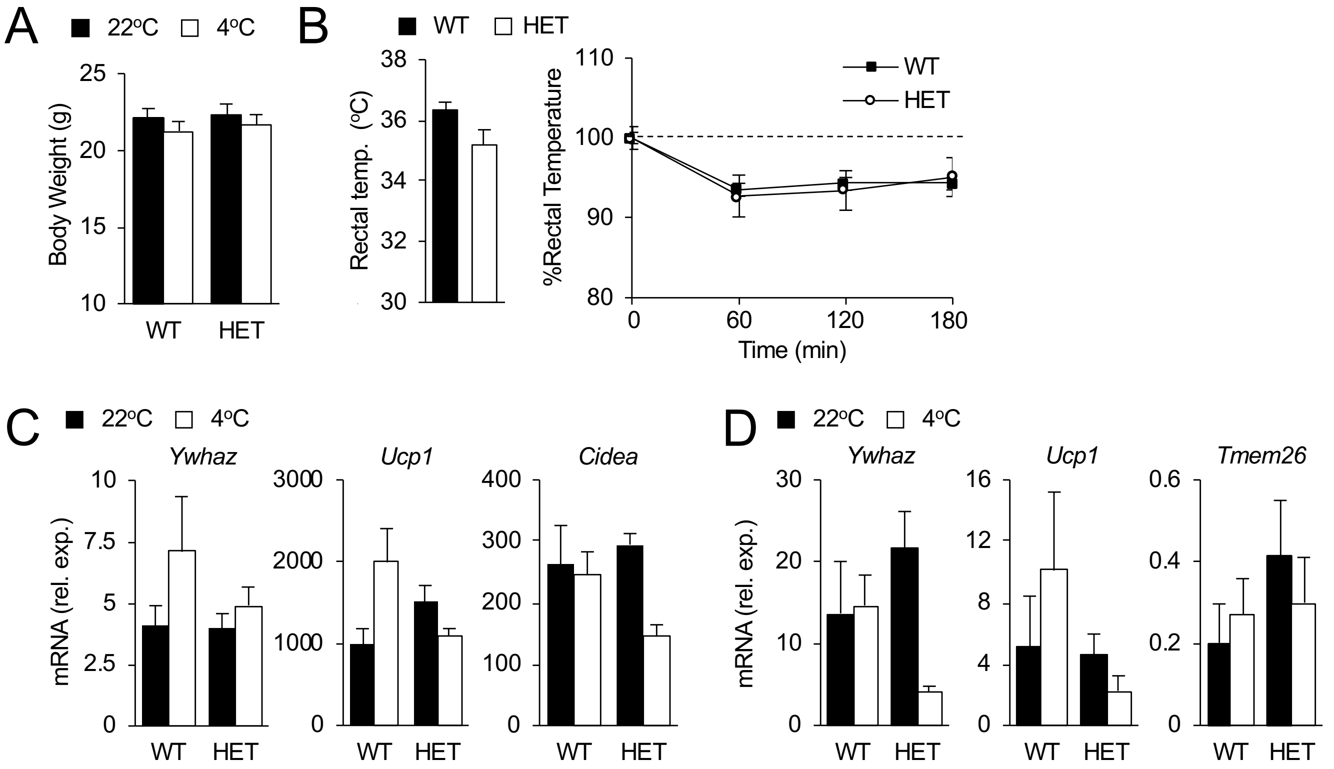


**Figure S1- Reducing 14-3-3ζ expression in female mice does not impact acute cold tolerance (A, B)** Body weights (B) and rectal temperatures (C) of female WT and HET mice were obtained prior, during, and at the end of the 3 hour cold challenge (n=7 mice per group). **(C,D)** Expression of brown-selective (C) and beige-selective (D) genes from BAT and iWAT, respectively, at room temperature (22 °C) and after 3 hours cold exposure (n=7 per group, *: $p<0.05$ when compared to 22 °C).

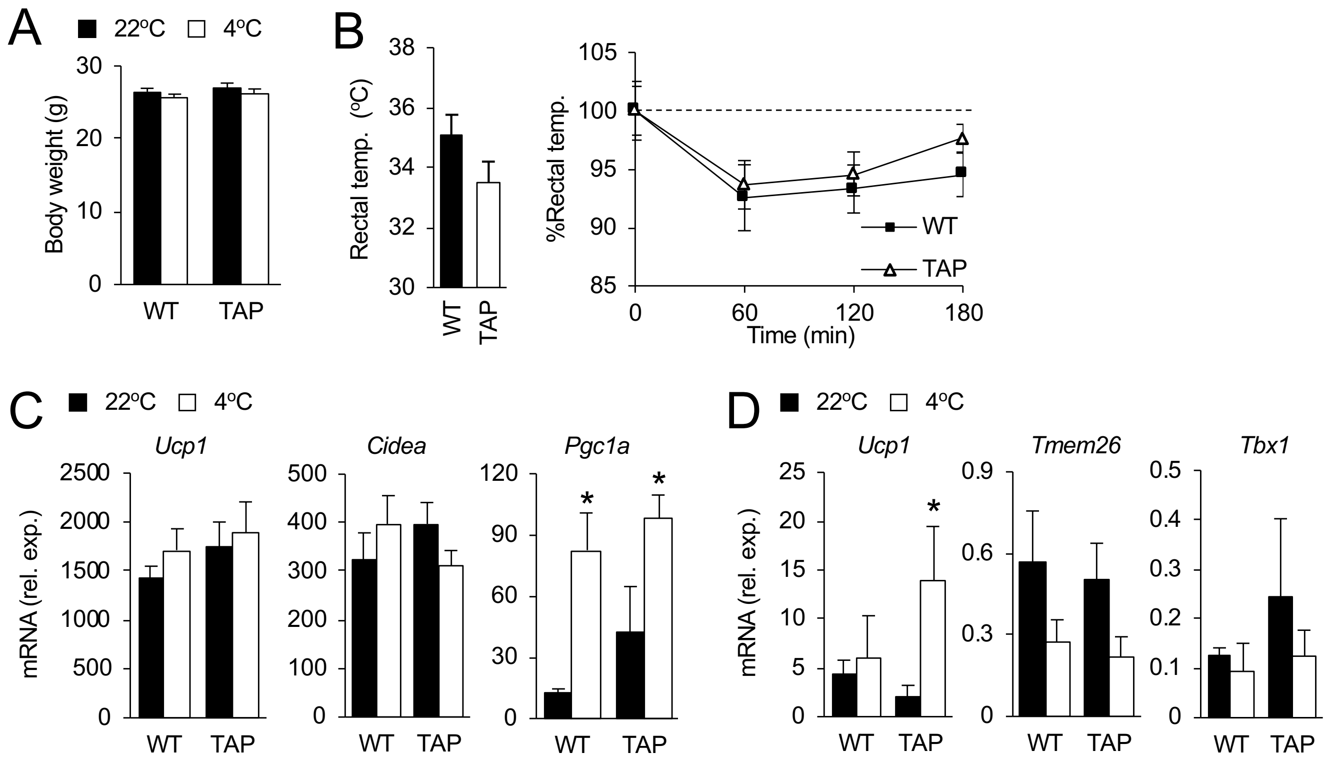


**Figure S2- Over-expression of 14-3-3ζ in female mice does not impact acute cold tolerance**
**(A,B)** Body weights (A) and rectal temperatures (F) of WT and TAP mice were obtained prior, during, and at the end of the 3 hour cold challenge (n=7 mice per group). **(C,D)** Expression of brown-selective (C) and beige-selective (D) genes from BAT and iWAT, respectively, at room temperature (22 ºC) and after 3 hours cold exposure (n=7 per group, *: $p<0.05$ when compared to 22 ºC).

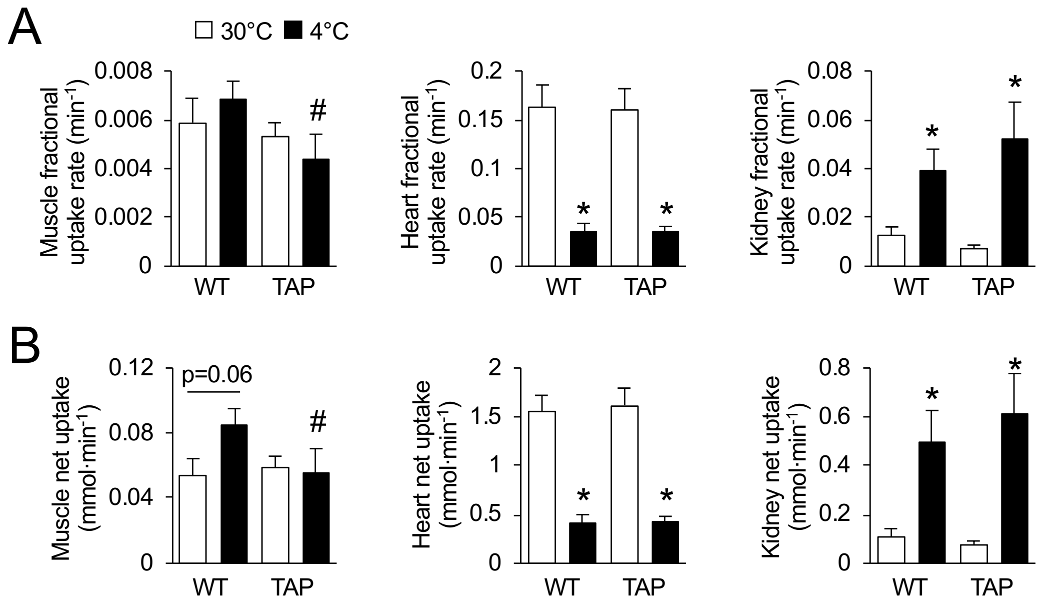


**Figure S3**: **14-3-3ζ over-expression does not alter [$^{18}$F]-FDG uptake in the heart or kidney of cold exposed mice.** Following exposure to thermoneutrality (30°C) or 4°C for 3 days, Fractional (A) and net (B) [$^{18}$F]-FDG uptake was measured in muscle, heart, and kidneys from WT and TAP mice at the indicated temperatures (n=8 WT, 10 TAP; *: p<0.05).

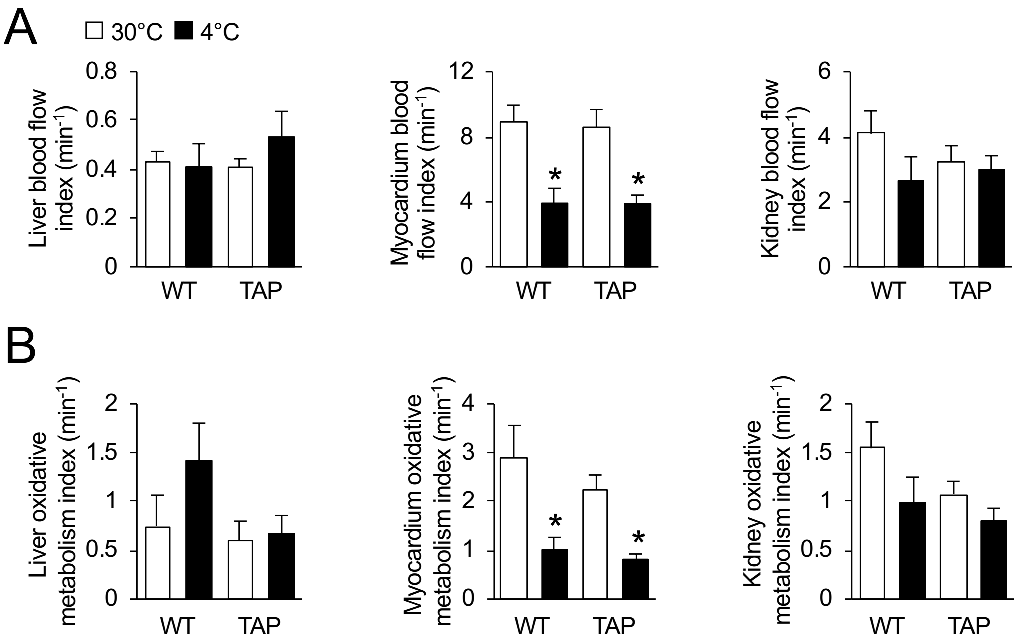


**Figure S4: No differences in oxidative metabolism or blood flow were detected in non-adipose tissues following cold exposure.** After being housed at thermoneutrality (30°C) or 4°C for 3 days, blood flow (A) and oxidative activity, as measured by [$^{11}$C]-acetate metabolism (B), were measured in myocardium, liver, and kidneys of WT and TAP mice at the indicated temperatures (n=8 WT, 10 TAP; *: $p<0.05$).

**Table S1: List of primers for qPCR**

| Gene | Forward | Reverse | Ref. |
|---|---|---|---|
| Adrb3 | CCTTCAACCCGGTCATCTAC | GAAGATGGGGATCAAGCAAGC | 1 |
| Cidea | TGCTCTTCTGTATCGCCCAGT | GCCGTGTTAAGGAATCTGCTG | 2 |
| Fgf21 | CTGGGGGTCTACCAAGCATA | CACCCAGGATTTGAATGACC | 3 |
| Hprt1 | TCCTCCTCAGACCGCTTTT | CCTGGTTCATCATCGCTAATC | 4 |
| Pdk4 | CCGCTTAGTGAACACTCCTTC | TCTACAAACTCTGACAGGGCTTT | 2 |
| Pgc1a | AGCCGTGACCACTGACAACGAG | GCTGCATGGTTCTGAGTGCTAAG | 2 |
| Pparg2 | GTTATGGGTGAAACTCTGGGAGAT | GGCCAGAATGGCATCTCTGTGTCAA | 4 |
| Prdm16 | CAGCACGGTGAAGCCATTC | GCGTGCATCCGCTTGTG | 2 |
| Tbx1 | GGCAGGCAGACGAATGTTC | TTGTCATCTACGGGCACAAAG | 2 |
| Tmem26 | ACCCTGTCATCCCACAGAG | TGTTTGGTGGAGTCCTAAGGTC | 2 |
| Ucp1 | ACTGCCACACCTCCAGTCATT | CTTTGCCTCACTCAGGATTGG | 2 |
| Ywhaz | CAGAAGACGGAAGGTGCTGAGA | CTTTCTGGTTGCGAAGCATTGGG | 5 |
| YWHAZ | ACCGTTACTTGGCTGAGGTTGC | CCCAGTCTGATAGGATGTGTTGG | 5 |


1. Lee Y, Petkova AP, Konkar AA, Granneman JG. Cellular origins of cold-induced brown adipocytes in adult mice. FASEB J, 29(1) (2015) 286-299.

2. Wu J, Boström P, Sparks LM, Ye L, Choi JH, Giang AH, Khandekar M, Virtanen KA, Nuutila P, Schaart G, Huang K, Tu H, van Marken Lichtenbelt WD, Hoeks J, Enerbäck S, Schrauwen P, Spiegelman BM. Beige adipocytes are a distinct type of thermogenic fat cell in mouse and human. Cell, 150(2) (2012) 366-76.

3. Jornayvaz FR, Birkenfeld AL, Jurczak MJ, Kanda S, Guigni BA, Jiang DC, Zhang D, Lee HY, Samuel VT, Shulman GI. Hepatic insulin resistance in mice with hepatic over-expression of diacylglycerol acyltransferase 2. Proc Natl Acad Sci U S A., 108(14) (2011) 5748-52.

4. Mugabo Y, Sadeghi M, Fang NN, Mayor T, Lim GE. Elucidation of the 14-3-3ζ interactome reveals critical roles of RNA-splicing factors in adipogenesis. J Biol Chem, 293(18) 6736-50.

5. Lim GE, Piske M, Johnson JD. 14-3-3 proteins are essential signalling hubs for beta cell survival. Diabetologia, 56(4) (2013) 825-37.

**Table S2: List of Antibodies**

| | Compay | Dilution | Product number | RRID |
|---|---|---|---|---|
| Alexa Fluor 594 | Jackson Immunoresearch | 1:400 | 115-585-003 | AB_2338871 |
| Anti-Mouse HRP IgG | Cell signaling | 1:5000 | 7076S | AB_330924 |
| β-Actin | Cell signaling | 1:10000 | 3700S | AB_2242334 |
| β-Tubulin | Cell signaling | 1:10000 | 86298S | AB_2715541 |
| GAPDH | Cell signaling | 1:10000 | 5174S | AB_10622025 |
| Perilipin | Cell signaling | 1:400 | 9349S | AB_10829911 |
| TH | Millipore | 1:400 | MAB318 | AB_2201528 |
| TH | Millipore | 1:400 | AB152 | AB_390204 |
| UCP1 | ABCAM | 1:1000 | Ab10983 | AB_2241462 |
| UCP1 | R&D systems | 1:400 | MAB6158-SP | AB_10572490 |
